# Contribution of self- and other-regarding motives to (dis)honesty

**DOI:** 10.1101/590208

**Authors:** Anastasia Shuster, Dino J Levy

## Abstract

Why would people tell the truth when there is an obvious gain in lying and no risk of being caught? Previous work suggests the involvement of two motives, self-interest and regard for others. However, it remains unknown if these motives are related or independently contribute to (dis)honesty, and what are the neural instantiations of these motives. Using a modified Message Game task, in which a Sender sends a dishonest (yet profitable) or honest (less profitable) message to a Receiver, we found that these two motives contributed to dishonesty independently. Furthermore, the two motives involve distinct brain networks: the LPFC tracked potential value to self, whereas the rTPJ tracked potential losses to other, and individual differences in motives modulated these neural responses. Finally, activity in the vmPFC represented a balance of the two motives unique to each participant. Taken together, our results suggest that (dis)honest decisions incorporate at least two separate cognitive and neural processes – valuation of potential profits to self and valuation of potential harm to others.

## Introduction

Why do people lie so often, even though honesty is a social norm? Then again, why would they tell the truth, in the face of possible monetary gain and no chance of punishment? Previous research suggests that the decision to lie or tell the truth depends on (1) the size of the profit to oneself gained from lying (*self-interest*), and (2) the degree of harm that the lie would cause to others (*regard for others*)^1^. Other self-related motives, such as the chance of being caught^2^, maintaining a positive self-image^2,3^, and an aversion to lying^4^ also decrease dishonesty. Less research has looked into how *outcomes* of dishonest behaviour affect it^1^. Indeed, lies are often self-serving but at the same time they are also other-harming (though not always^5,6^). Moreover, the *degree* to which a given opportunity to lie is self-serving and other-harming varies from one scenario to the other. For example, the amount of money in a wallet found on the street entails both the potential profit for the finder and the loss for the owner. Recently, researchers found that the more money a wallet held, the *more* likely people were to surrender it to the police^7^. This supports the observation that people are sensitive to their own payoff as well as another’s loss when confronted with a moral choice. However, it is not clear whether the two pieces of information distinctly affect behaviour, and whether people vary in the degree to which either self- or other-regarding motives drive their behaviour. The neural computation underlying the arbitration between these conflicting motives remains elusive as well.

When studying deception, researchers often use so-called instructed lying paradigms^8,9^. Although they enable significant experimental control, they suffer from low ecological validity. In these tasks, participants do not benefit from lying (thus nullifying the conflict between moral values and self-interest). More importantly, lying in such scenarios has no social consequences, since the tasks are non-interactive and the lie does not harm anyone (thus nullifying the conflict between self-interest and regard for others’ wellbeing^9^). Recently, a group of researchers put forward a signalling framework drawing from game theory^10^ to study spontaneous dishonesty^8^ in socially interactive settings^11^, preserving the conflicts of real-life dishonesty. We used a modified version of one such task, the Message Game, in which a *Sender* sends either a truthful or a deceiving message to a *Receiver* regarding which of two options to choose. The task conflicts monetary gain to the Sender with honesty, to invoke internally-motivated lying. As potential profits to the Sender go up, Senders tend to send the deceptive message more often. Conversely, as potential losses to the Receivers rise, Senders lie less^1^. By systematically varying the payoffs to both players, we can estimate the role of self- and other-regarding motives in dishonest choices and track their neural correlates. In the original Message Game, Sender’s message does not always predict the Receiver’s choice^1,4^. This can lead to strategic choices, in which a Sender might choose to tell the truth while having the intention to deceive^12^. Therefore, we modified the task to include two options of $0 to both players. Thus, deviating from the Sender’s message would result in a 66% chance of not winning money at all. This modification ensures that Senders’ choices have true consequences for their partners – a central component in our study.

Neuroimaging studies consistently implicate several regions of the prefrontal cortex (PFC) in generating dishonest behaviour^11,13–18^. For example, deceptive responses—but not erroneous ones—selectively activate the middle frontal gyrus^14,19^. Purposefully withholding information from the experimenter engages the anterior cingulate cortex, dorsolateral prefrontal cortex (dlPFC), and inferior and superior frontal gyrus^20–23^. Because most studies employed paradigms with no interpersonal interaction^11^ and thus no real consequences on another person, the neural correlates of the *social* aspect of deception are less well understood. However, there is plentiful evidence for the integration of other-regarding motives with self-interest during decision-making, coming from studies of prosocial behaviour. Even in the absence of explicit extrinsic pressure, humans are willing to forego monetary gain to cooperate^24^, share resources^25–27^ and act fairly^28^. Prosocial behaviour activates areas of the neural valuation system, consisting of the ventromedial prefrontal cortex (vmPFC) and ventral striatum^26,28^ (for reviews see^29,30^). In these regions, social (e.g., norms) and non-social (e.g., monetary profit) factors are integrated into a common decision-value, which then gives rise to choice^31–33^. Putatively, when choices present a conflict between one’s own profit and normative social principles, the valuation system interacts with areas typically involved in social cognition (e.g., the temporoparietal junction (TPJ)^34^ or the posterior superior temporal sulcus (STS)^27^) to compute the subjective value of an alternative^30^. In the present study, we focus on the unique contribution of self- and other-regarding motives driving (dis)honest behaviour, as well as the combination of the two by the valuation system.

Participants played as Senders, choosing between honest and dishonest alternatives while inside the fMRI scanner. Choosing the dishonest option resulted in higher gains for themselves and greater losses to their partner compared to the honest alternative. Our first aim was to measure how increases to own profit and other’s loss affect dishonest choices. Our second aim was to explore individual differences in these motives. Third, we aimed to identify the neural activity associated with the two value parameters that drive dishonest behaviour – value to self and other. Finally, we examined how the neural representation of these value parameters reflects individual differences in behaviour.

## Results

### Behaviour

On each trial in the task, the participant (*Sender*) chose to send either a truthful or a deceptive message to the *Receiver* (see Fig. 1). We started with a simple measure of overall dishonesty – on how many of the trials did participants chose to send the deceptive message. The deceptive message was sent on almost half of the trials, but with substantial variability between participants (45.45%±17.7%; range: 17.36%-89.93%). Overall dishonesty did not differ statistically between female and male participants (females: 43%±15.9% males: 50.59%±21%; *t*(27)=-1.06, *p*=0.29, two-tailed two-sample t-test). Participants took on average 2.87s to choose, and average reaction times for Truth choices (2.86±0.45 s) did not differ from reaction times for Lie choices (2.88±0.55 s; t(27)=0.4, *p*=0.69). The difference between honest and dishonest decision times, however, was significantly related to the individual differences in overall dishonesty: participants who lied more often took longer to tell the truth, whereas more honest participants took longer to lie (r(26)=-0.8, *p*<0.0001; Figure 2a). This suggests that the decision process reflects individual differences in dishonesty preference, not only in which alternative is chosen but also in the time it takes to choose.

**Figure 1.**
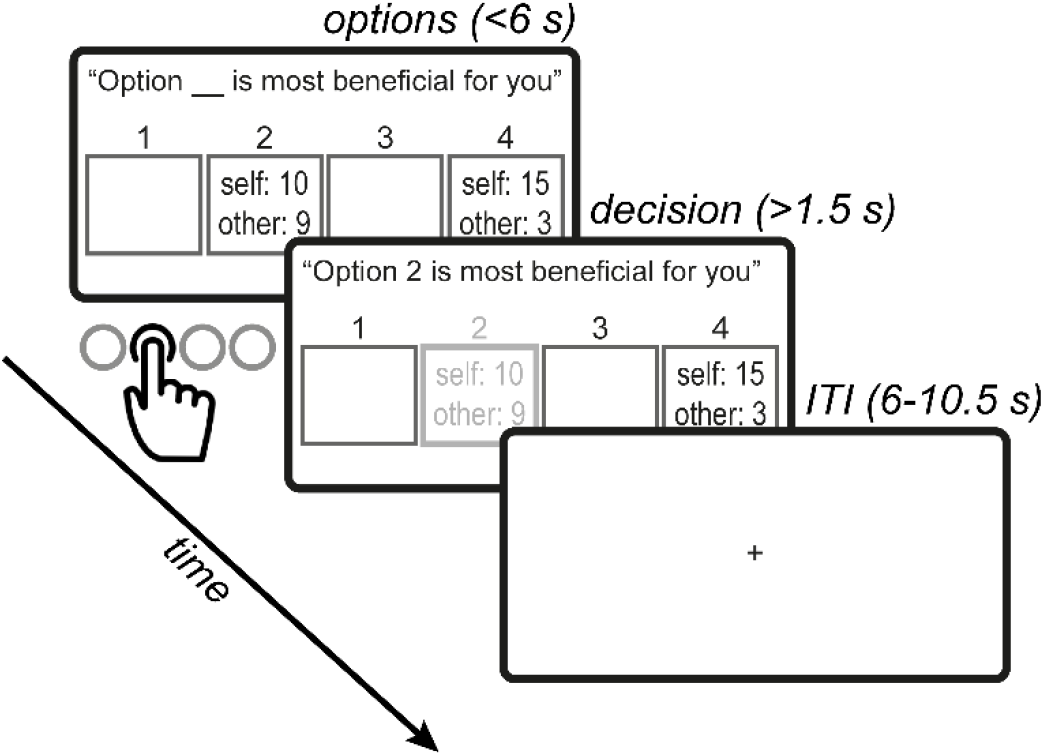
Trial timeline. On each trial, the participant (a “Sender”) chose which message to send to the Receiver, out of four options. The text at the top of the screen is the message that would be sent on this trial to the Receiver. Four options are revealed, each one consisting of some amount of money for the Sender (“self”) and some for the Receiver (“other”). One option was always truthful (the one more beneficial for the Receiver; #2 in the example) and one deceptive (#4). Payoffs to both players and locations varied between trials. The Sender had 6 seconds to indicate her choice, after which the chosen option was highlighted and stayed on the screen for the remainder of the trial.

**Figure 2.**
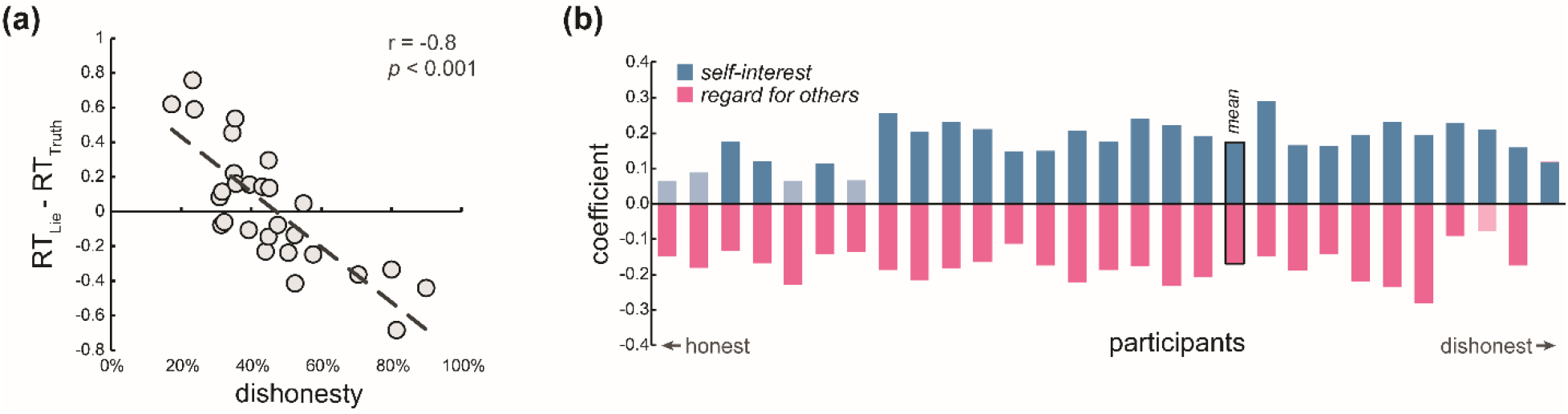
Behavioural results (n=28). **(a)** Reaction times correlate with overall dishonesty. The average difference between Lie reaction times and Truth reaction times are on the y-axis, and percentage of dishonest trials on the x-axis. Each circle represents a subject. **(a)** Participants are arranged from most honest (far left) to most dishonest (far right). Bars represent each participant’s regression coefficients, reflecting how much self-interest and regard for others contributed to their probability to lie. Greyed-out bars indicate non-significant coefficients.

#### Motives for (dis)honesty

To uncover what drove dishonest behaviour, we conducted a regression analysis of the probability to lie as a function of the potential payoffs, separately for each participant. The regression revealed that both the potential profits for the Sender (*value to self*; *ΔV*_self_) and the potential losses to the Receiver (*value to other*; *ΔV*_other_) affected the behaviour of most participants (Figure 2b). A large *ΔV*_self_ coefficient implies a more *self-interested* participant: each unit of money to the Sender causes a bigger increase in the probability to lie, compared to a small *ΔV*_self_ coefficient. In other words, self-interest represents the sensitivity of a person to their own profit when faced with temptation. Note, that a high *ΔV*_self_ coefficient (more sensitive to own profit) does not imply that this person is necessarily more dishonest overall (which is given by the regression’s intercept), but rather that when considering whether to lie or not, her own profit plays a more significant role in the decision. Similarly, a large *ΔV*_other_ coefficient (high *regard for others*) means that the loss to the Receiver greatly decreases the probability of the Sender to lie, compared to a small coefficient. While the average contribution of both parameters was similar in absolute terms (β*ΔV*_self_: 0.174±0.06, β*ΔV*_other_: *M*=-0.169±0.057, t(27)=0.35, *p*=0.73, paired two-tailed t-test), they varied substantially between participants (β*ΔV*_self_ range: 0.06-0.29; β*ΔV*_other_ range: −0.28-0.004). The fact that both coefficients were significant for the majority of participants implies that self-interest and regard for others independently affect the probability of the Sender to lie. This independence is further strengthened by a lack of correlation between the two coefficients across participants (r(26)=-0.2, *p*=0.3; Supplementary Figure S2).

### Neuroimaging

#### Neural correlates of dishonesty

To identify neural correlates of dishonest behaviour, we contrasted the neural response during trials in which participants lied (i.e., sent the deceptive message) with trials in which they told the truth, controlling for reward amount. Consistent with previous studies of deception^11,15^, we found several regions, including the medial PFC, left dlPFC, and bilateral insula. The opposite comparison (Truth>Lie) revealed activations in the right TPJ, right STS and cerebellum (see Table 1 and Supplementary Fig. S3).

**Table 1.** List of whole-brain contrast results. PFC: prefrontal cortex; IPL: inferior parietal cortex; TPJ: temporoparietal junction; SMA: supplementary motor area.

| contrast | brain area | MNI coordinate | | | $T$ | cluster size |
| --- | --- | --- | --- | --- | --- | --- |
| | | $x$ | $y$ | $z$ | | |
| *Truth vs. Lie* | medial PFC | -3 | 29 | 38 | 4.06 | 128 |
|  | (left) dorsolateral PFC | -47 | 31 | 28 | 3.93 | 23 |
|  | (left) insula | -34 | 23 | -2 | 4 | 31 |
|  | (right) insula | 30 | 25 | -2 | 4.37 | 29 |
|  | (right) TPJ | 64 | -28 | -25 | -4.02 | 116 |
|  | (left) superior temporal sulcus | -58 | -18 | -5 | -4.09 | 43 |
|  | cerebellum | -9 | -65 | -5 | -4.44 | 81 |
| *value to self*<br>($\Delta V_{self}$) | (left) lateral PFC | -44 | 16 | 31 | -3.53 | 409 |
|  | (right) IPL | 36 | -50 | 47 | -3.62 | 474 |
|  | (left) IPL | -26 | -54 | 39 | -3.6 | 1471 |
|  | pre-SMA | -2 | 15 | 50 | -3.42 | 51 |
|  | (left) occipital | -45 | -67 | 4 | -3.33 | 72 |
|  | (right) occipital | 45 | -77 | -3 | -3.4 | 168 |
|  | (medial) occipital | -15 | -70 | 1 | -3.57 | 666 |
|  | cerebellum | 5 | -64 | -39 | -3.35 | 63 |
|  | (right) amygdala | 28 | -4 | -14 | -3.5 | 46 |
| *value to other*<br>($\Delta V_{other}$) | (right) TPJ | 45 | -52 | 24 | -3.37 | 33 |
|  | precuneus | -7 | -63 | 44 | 3.79 | 21 |
|  | (left) thalamus | -6 | -18 | 11 | 3.37 | 26 |
|  | cuneus | -6 | -92 | 22 | 3.36 | 32 |
|  | (left) occipital | -9 | -73 | 3 | -3.4 | 115 |
|  | (right) occipital | 9 | -73 | -8 | -3.42 | 108 |

#### Chosen value

To identify voxels responding to the Sender’s reward magnitude in the chosen alternative vs. the unchosen one, we parametrically modelled the Sender’s expected reward based on her choice and looked within the value system^35^. Consistent with previous findings^35,36^, the amount of money in the chosen vs. the unchosen option for the Sender positively correlated with the BOLD signal in the valuation system – the vmPFC and bilateral ventral striatum (see Fig. 3a).

**Figure 3.**
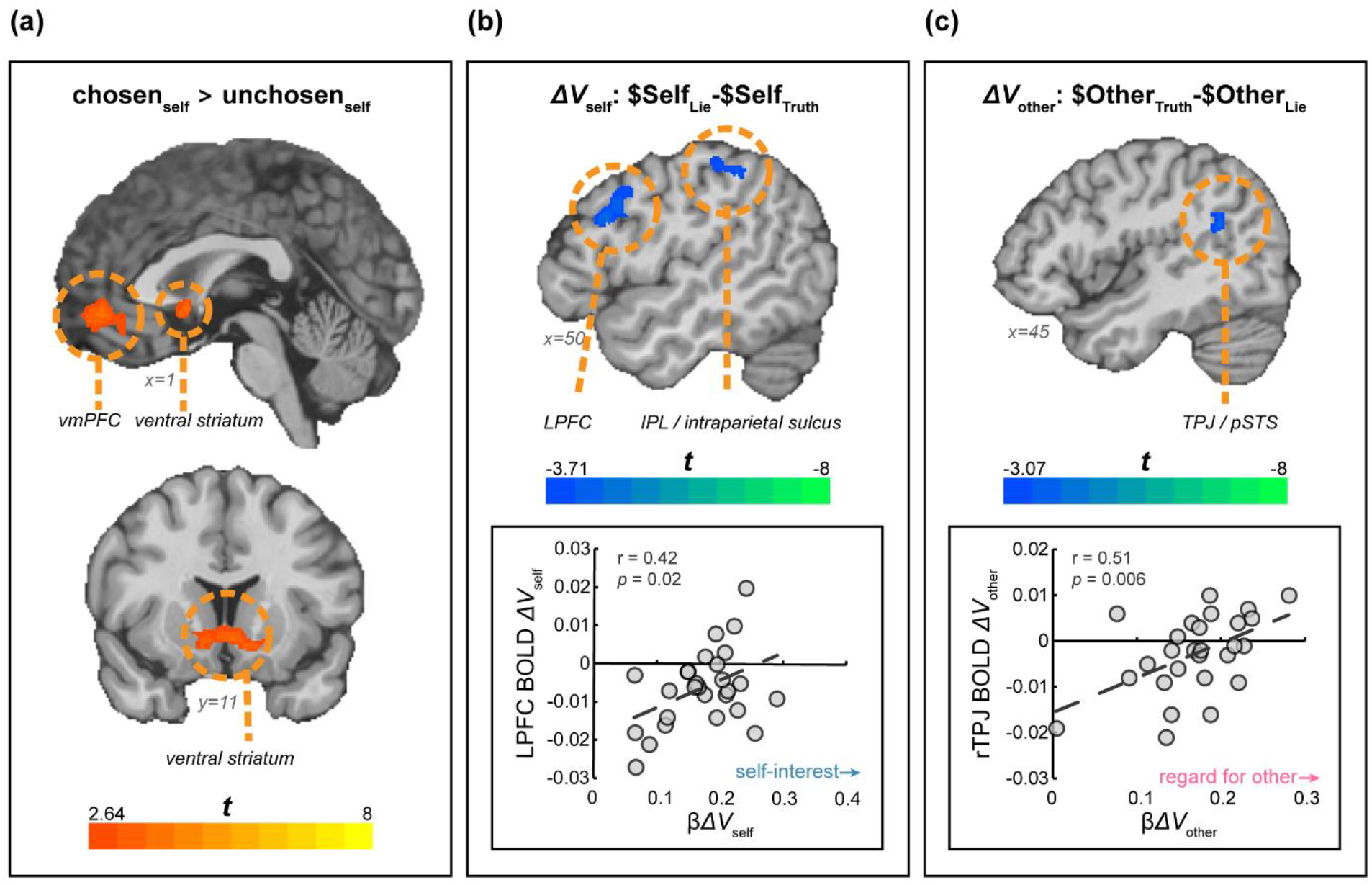
Neural sensitivity to values (n=27). **(a)** Amount of money for the Sender in the chosen option vs. the unchosen option. The activation map was masked using value-related ROIs taken from a meta-analysis (Bartra et al., 2013). Map at *q*(FDR)=0.05. **(b)** Voxels sensitive to Sender’s potential profits from dishonesty (value for self; *ΔV*_self_) (top). For visualization purposes, the map is thresholded at *p*=0.001, cluster-size corrected. The BOLD response in the LPFC is positively correlated with the behavioural measure of self-interest (bottom). **(c)** Voxels sensitive to Receiver’s potential losses from dishonesty (value for other; *ΔV*_other_) (top). Map thresholded at *p*=0.005, cluster-size corrected. The BOLD response in the rTPJ is positively correlated with the behavioural measure of regard for others.

#### Value for self and other

##### Value to Self

We found that the left lateral prefrontal cortex (LPFC) and intraparietal lobule, among other regions, negatively tracked the amount of money a Sender can gain from lying (*ΔV*_self_; Table 1, Figure 3b). That is, a smaller potential profit from lying corresponded to a higher activation in the left LPFC and IPL. To examine how individual differences in self-interest affect this neural activity, we extracted the BOLD coefficients for *ΔV*_self_ from an independently defined region of interest (ROI) of the LPFC (MNI coordinates x, y, z: −48, 6, 28^37^) and correlated them with β*ΔV*_self_ estimated using participants’ behaviour. We found a positive significant neural-behaviour correlation (r(25)=0.42, *p*=0.027), where less self-interested participants show more deactivations of the LPFC.

##### Value to Other

To identify voxels tracking the value the Sender exerts toward the Receiver (i.e., value to other, *ΔV*_other_), we regressed the potential monetary loss to the Receiver if the Sender chooses to act dishonestly ($Other_Lie_-$Other_Truth_). We found an inverse relationship between the activity of the rTPJ and Receiver’s potential loss, such that higher potential losses to the Receiver deactivated the rTPJ (Table 1; Figure 3c). We further extracted neural responses from the rTPJ using an independently-defined ROI (MNI coordinates x, y, z: 54, −58, 22^37,38^) and examined whether differences between participants in the activity of the rTPJ can be explained by individual differences in their other-regarding motive for lying. We found that participants with low regard for others showed rTPJ deactivation, whereas in those who have high regard for others, higher potential losses elicit *more* activity in the rTPJ (r(25)=0.51, *p*=0.006). That is, the degree to which social consequences affect an individual’s behaviour is related to how these social consequences are represented in the rTPJ.

#### Balanced self-other representation

To understand how the vmPFC tracks competing values and motives, we looked for differences between neural sensitivity to value to self and value to other in the valuation system, and how self- and other-regarding motives affect this activity. To do so, we first computed for each participant an *other-self differential* score (β*ΔV*_other_–β*ΔV*_self_), indicating how much a participant’s behaviour is driven by one motive or the other. High positive scores indicate a higher contribution of other-regarding motives, whereas high negative scores suggest more contribution of self-regarding motives. Then, we compared the neural activity tracking value to other with that of value to self (*ΔV*_other_ > *ΔV*_self_) in the valuation system. This contrast yielded an empty map. However, when we regressed onto this map each participant’s *other-self differential* score, we identified a cluster of voxels located in the vmPFC (MNI coordinate x, y, z: 9, 41, −8; Figure 4). The vmPFC was more active for *ΔV*_self_ compared to *ΔV*_other_ in self-interested participants. Conversely, for other-regarding participants, who place higher weight on *ΔV*_other_ compared to *ΔV*_self_, the vmPFC was more active for *ΔV*_other_ compared to *ΔV*_self_ (r(25)=0.49, *p*=0.009). Thus, the vmPFC represents in each participant her own *idiosyncratic balance* between value to self and value to other. This finding is consistent with the role of the vmPFC as an integrator of various attributes of choice to one single value.

**Figure 4.**
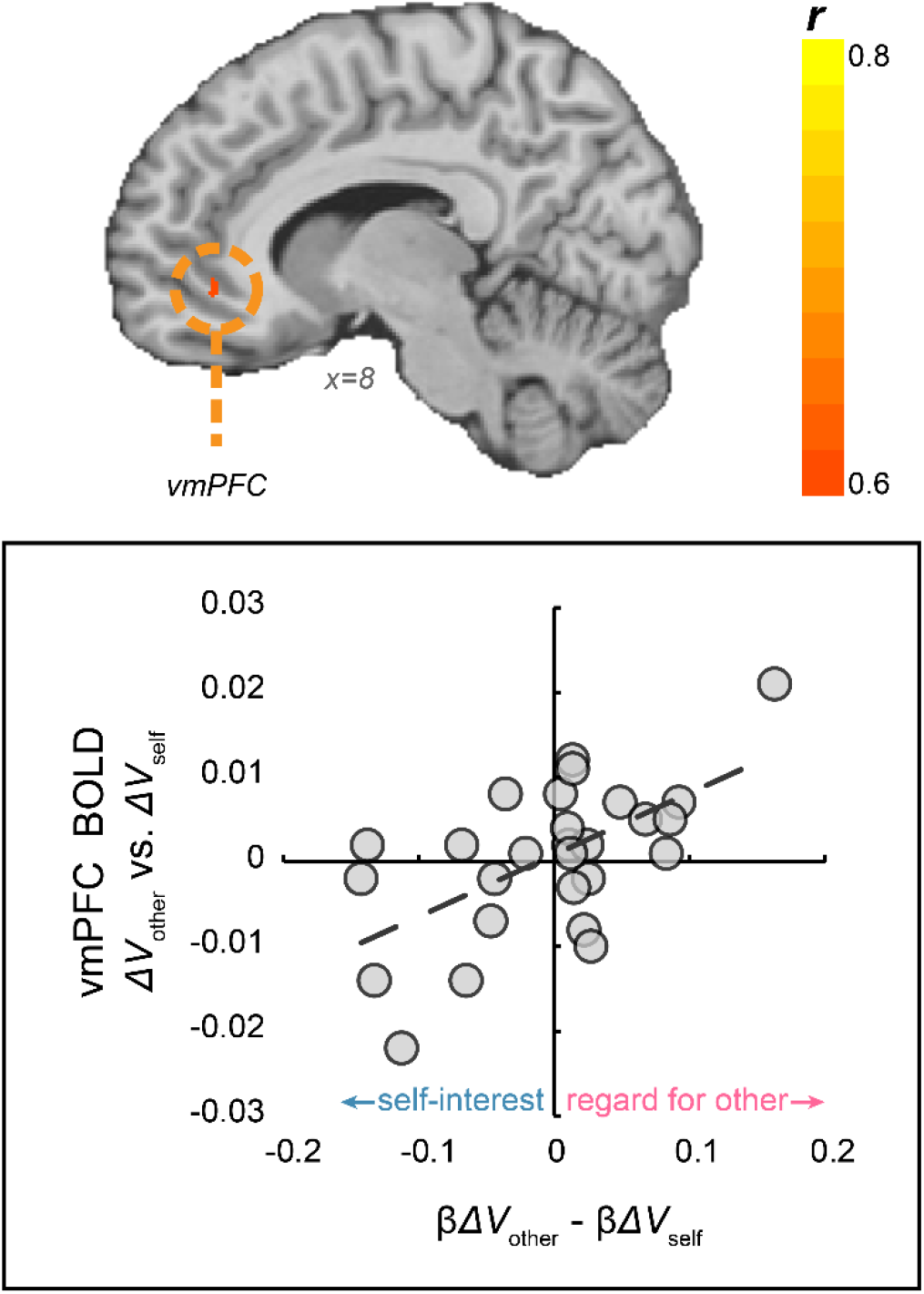
Value to self vs. value to other, modulated by preferences (N=27). Contrasting potential profit for Sender (value to self; *ΔV*_self_) with potential losses to Receiver (value to other; *ΔV*_other_), modulated by individual differences in the balance between regard for other and self-interest. The activation map was masked using value-related ROIs taken from a meta-analysis (Bartra et al., 2013).

## Discussion

Using a modified Message task, we found that both self-interest and regard for others contribute independently to (dis)honesty. Crucially, our results suggest that these motives rely on distinct neural processes, with self-interest involving the lateral prefrontal and parietal cortices, and regard for others the right temporoparietal junction, among others. Furthermore, we find a combination of motives in the vmPFC, consistent with its known role as an integrator of value. That is, in self-interested participants, the vmPFC was more sensitive to one’s own payoffs than losses to another; conversely, in other-regarding participants, the vmPFC showed higher activity for their partner’s potential losses than to their own potential gains.

Previous research has indicated that increasing the consequences of the lie decreases its occurrence^1^. We have extended this notion to elucidate individual differences in this sensitivity, and thanks to our within-participant design and orthogonality of the regressors, we captured independent self- and other-regarding motives. This independence coincides with the observation that some people would not lie, even if lying would *help* another person (as in a case of a white lie^39^), suggesting that for some people, the self-regarding motive is the *only* motive driving behaviour and that honesty can be wholly unrelated to the consequence of lying.

In the context of choices, differences in reaction times should be interpreted carefully. Reaction times could attest to the ease of choice^40,41^, or to the rate in which evidence accumulates in favour of one of the alternatives^42^, which is related to the strength-of-preferences^41^. Although all participants were faced with the same choice set, variation in their preferences yielded variations in decision times. Along the same lines as previous findings^41^, we find that honest participants are quicker to tell the truth, whereas dishonest participants are quicker to lie. In other words, acting out of character takes longer. This result goes against dual-process models, which assume similar preferences for all participants (e.g., everyone is honest at heart, and lying requires extra time to overcome the honesty urge).

We find that potential profits from lying involve the LPFC, dovetailing some recent research elucidating its role in moral decision-making^43^. Lesions to the dlPFC were found to reduce honesty concerns in favour of self-interest^44^, suggesting it is causally involved in arbitrating the two. Further evidence comes from a stimulation study^45^, in which activating the dlPFC increased honesty. Interestingly, this was only true when a conflict existed between self-interest and honesty, suggesting that, as the authors describe it, the role of the dlPFC is to represent the psychological cost of being dishonest. The LPFC was also implicated in another type of moral decision-making, when the harm to another individual is physical (electric shocks) rather than monetary^37^. Not only did the LPFC negatively represent potential ill-gotten gains, but individual differences in harm-aversion correlated with the activity in the LPFC. Specifically, activity in this region decreased as the amount of money that could be gained grew, paralleling our own findings in a different task and another moral domain. The authors attributed this finding to a sense of blame, showing that the LPFC encodes the level of blameworthiness – higher gains are associated with lower blame and hence lower activity in the LPFC, echoing the notion of psychological cost. Taken together, these studies, along with our own, support the role of the LPFC as a context-specific modulator of decision-relevant information^43^.

As for the other-regarding motive to act honestly, we find that the right TPJ negatively tracks the value for other (*ΔV*_other_). We further find that the rTPJ encodes the Receiver’s loss positively in participants who care more about it. Thus, the sensitivity of the rTPJ to consequences of dishonesty can drive some individuals to act prosocially and others selfishly. Interestingly, although this area has been implicated in deception, it does not show up in the overlap between deception-related and executive-control-related brain maps^15^, suggesting that it may represent a different aspect of dishonesty. We propose that this aspect is its social consequences. Our hypothesis is supported by a meta-analysis of deception studies, showing that the TPJ is specifically involved in interactive deception tasks^11^, that is, only when a lie has implications on another participant. Indeed, the TPJ is a known hub in the social cognition neural network, selectively activated when interacting with social agents^46^, and is thought to represent others’ minds^47–49^. The locus of our activation corresponds to the posterior rTPJ, an area highly connected with the vmPFC and implicated in a wide array of mentalizing tasks^49^. In the moral domain, the rTPJ is involved in forming judgments about others’ moral acts^50,51^. Here, we provide evidence that the potential *outcomes of moral acts* also involve the rTPJ. Moreover, we show that the individual’s neural sensitivity to these potential outcomes can explain their social behaviour.

Finally, we find that the vmPFC represents a balance between competing values. Importantly, this balance in the vmPFC emerges only when accounting for the weight that each individual placed on their own and another’s payoff, implying that the nature of this representation is subjective – personal preferences modulate the objective amounts of money. We find that if someone cares more about themselves than others, her valuation system will be more sensitive to her own profits. Alternatively, the valuation system of someone who is concerned with others more than with herself, would be more sensitive to others’ profits. This balanced representation of value suggests a neural instantiation of observed differences in behaviour. Our findings are in line with abundant research on the role of the vmPFC in value computation and representation^27,32,33,46^, suggesting that when two values or motives conflict, the vmPFC represents a comparison of them.

In summary, we found that internally-motivated dishonest behaviour^9^ varies dramatically between individuals, both in which motive drives behaviour and to what extent. Moreover, we find that two distinct motives can be identified from behaviour and traced back to separate neural activation patterns. At the behavioural level, individual’s levels of self-interest and regard for others contribute independently to a choice to lie. On the neural level, the LPFC and rTPJ track value to self and value to other, respectively. Importantly, self- and other-regarding motives for (dis)honesty affect this sensitivity to value – in the LPFC, the rTPJ, and the valuation system. These findings suggest a neural instantiation of individual differences in behaviour; while prosocial individuals’ neural valuation systems are more sensitive to others’ wellbeing, those of selfish individuals are more sensitive to their own.

## Materials and Methods

### Participants

Thirty-three participants enrolled in the study (22 females; age *M*=25.35, 19-30). Participants gave informed written consent before participating in the study. All experimental protocols were approved by the Institutional Review Board at Tel-Aviv University, and all methods were carried out in accordance with the approved guidelines. All participants were right-handed, and had a normal or corrected-to-normal vision. Of the 33 participants, five were excluded from all analyses due to a lack of variability in their behavioural responses – they either lied on more than 90% (n=3) or on less than 10% (n=2) of the trials. An additional participant was excluded only from neural analyses due to excessive head movements during the scans (>3 mm). Participants were paid a participation fee and the amount of money they won on a randomly drawn and implemented trial.

### Experiment

#### Task

On each trial, the participant in the fMRI scanner (*Sender*) was asked to send the following message to her partner (*Receiver*): “Option ____ is most profitable for you”. Senders watched a screen with 4 ‘doors’. Two doors were empty (indicating payoffs of $0 for both participants), and two doors contained non-zero payoff information for themselves and for their partner (the Receiver; see Fig. 1). Participants were given 6 seconds to indicate their choice by pressing one of four buttons on a response box. After choosing, the chosen door was highlighted and the message text changed accordingly (e.g., “Option 3 is most profitable for you”, if door 3 was chosen). The duration of this decision screen was set to have a minimum of 1.5 s and to make up for a total trial duration of 7.5 s. For example, if a participant made a choice after 3.5 seconds, the decision screen appeared for 4 seconds. If no choice was made in the allotted time, a “no choice” feedback screen appeared for 1.5 s. Afterwards, a fixation screen appeared for 6-10.5 seconds. Senders were told that the Receiver would view the message and choose which door to open, when the only available information for them is the door number, and the fact that two of the doors are empty. Both players’ payoffs depended on the door the Receiver opened. To avoid reputation concerns, even after choice, the Receiver did not know how much the Sender got, nor what amounts of money were hidden behind the non-chosen doors.

#### Payoff structure

The payoffs were intended to create a conflict between honesty and monetary profit. On each trial, each door contained some amount of money for the Sender and some amount of money for the Receiver. A truthful message (*Truth* option) is defined as when the Sender is choosing the door (the message to send) that results in a larger amount of money for the Receiver. A deceptive message (*Lie* option) is defined as when the Sender is choosing the door that results in a smaller amount of money for the Receiver (and a larger amount of money for the Sender) compared to the Truth option. Payoffs varied on a trial-by-trial basis, ranging between 10-42₪ (1₪ ≈ 0.3USD) for the Sender, and 1-31₪ for the Receiver (see Supplementary Table S1 and Supplementary Fig. S1 for a full list). Critically, the potential profits for the Sender from lying (*ΔV*_self_, $Self_Lie_-$Self_Truth_; range 0-12₪) varied independently from the potential losses for the Receiver (*ΔV*_other_, $Other_Truth_-$Other_Lie_; range 1-20₪; r(38)=0.25, *p*=0.11). Figure 1 depicts an example trial, in which choosing to tell the truth would result in 10₪ for the Sender and 9₪ for the Receiver, whereas lying would result in 15₪ for the Sender and only 3₪ for the Receiver. Hence, the Sender’s potential profit (*ΔV*_self_) is 5₪ (15 − 10), and the Receiver’s potential loss (*ΔV*_other_) is 6₪ (9 − 3). Importantly, to avoid any interfering motives (e.g. envy), all trials of interest (38 out of 40) had a higher payoff for the Sender than for the Receiver (the Sender was always better off than the Receiver, irrespective of the chosen door). Two *catch trials* offered a 0₪ payoff for the Sender from lying (i.e., $Self_Lie_=$Self_Truth_), with no conflict between honesty and gain for the Sender. These trials served to ensure the participants are attentive.

A key component of the task is the addition of two empty doors to each trial, containing zero money for each player (doors 1 & 3 in the Fig. 1 example). These options ensured that the message the Sender sends is indeed followed by the Receiver, because deviating from the recommended door may result in opening an empty door. Consider the example in Fig. 1: if the Sender chooses to lie, she sends a message regarding door #4. All the Receiver will see is that door #4 is highlighted. The Receiver does not know how much money is behind which door, but she does know that two of the doors have no money at all. Even if the Receiver believes she is lied to, it is in her best interest to open door #4, otherwise she (and the Sender) face a 66% chance of winning no money at all.

#### Procedure

Participants arrived to the Imaging Center and met the experimenter and a confederate acting as the Receiver. The confederate was always a Caucasian female, aged ~25. Both the Sender and Receiver read written instructions, signed consent forms and underwent a training stage of the task – playing as both the Sender and the Receiver. Training as Sender was intended to familiarize the participant with the Message Task. Training on the Receiver’s role allowed them to experience what happens when the Receiver does not follow the recommended Message, and ensure they understood the consequences of sending a truthful or deceptive message.

Each participant completed four scans. Forty unique payoff trials were randomly interspersed in a given scan, making up 160 trials per participant (40 unique payoffs × 4 repetitions). At the end of the experiment, one trial was selected randomly and presented to the Sender as the message that will be sent to the Receiver. In pilot studies in our lab, using real participants acting as Receivers, 100% chose according to the message sent by the Sender. Therefore, in the current study, we automatically set the Receiver’s choice to be according to the Sender’s message, and paid the Sender whichever amount was associated with that option.

After the scan, participants completed a short debriefing questionnaire. Debriefing consisted of several demographic questions and questions regarding the choices the participant made in the task. Specifically, we aimed to ensure participants were not suspicious of the confederate, by asking how the identity of the Receiver affected their choices.

### Image acquisition and processing

Scanning was performed at the Strauss Neuroimaging Center at Tel Aviv University, using a 3T Siemens Prisma scanner with a 64-channel Siemens head coil. Anatomical images were acquired using MPRAGE, which comprised 208 1-mm thick axial slices at an orientation of −30° to the AC–PC plane. To measure blood oxygen level-dependent (BOLD) changes in brain activity task performance, a T2*-weighted functional multi-band EPI pulse sequence was used (TR = 1.5s; TE = 30 ms; flip angle = 70°; °; matrix = 86 × 86; field of view (FOV) = 215 mm; slice thickness = 2.5 mm). 50 axial (−30° tilt) slices with no inter-slice gap were acquired in ascending interleaved order.

BrainVoyager QX (Brain Innovation) was used for image analysis, with additional analyses performed in Matlab. Functional images were sinc-interpolated in time to adjust for staggered slice acquisition, corrected for any head movement by realigning all volumes to the first volume of the scanning session using six-parameter rigid body transformations, and de-trended and high-pass filtered to remove low-frequency drift in the fMRI signal. Data were then spatially smoothed with a Gaussian kernel of 5 mm (full-width at half-maximum), co-registered with each participant’s high-resolution anatomical scan and normalized using the Montreal Neurological Institute (MNI) template. All spatial transformations of the functional data used trilinear interpolation.

### Behavioural analysis

#### Overall deception & reaction times

First, we removed no-response trials (0.9%±1.2% of trials), and trials in which participants chose an empty door (option of $0 to both players; 1.4%±1.3% of trials). Finally, we removed the catch trials (trials with identical payoffs to the Sender for Truth and Lie; 4.8%±0.03% of trials). All further analyses were performed on this subset of trials (92.7%±3.2% of trials, range: 137-152 trials). We defined overall dishonesty rates per participant as the number of trials they lied out of this total number of trials.

#### Analysis of motives

To estimate the contribution of self- and other-regarding motives to dishonest behaviour, we fitted a linear regression per participant. Note, that there is increased variability in value-to-self compared to value-to-other. This was done for experimental reasons, to elicit sufficient lying. To allow us to still compare the regression coefficient across variables, we have z-transformed both variables. Then, for each unique payoff, we calculated each participant’s probability to lie by averaging choices across the four repetitions, and that served as the dependent variable. The independent variables were normalized profits to self and losses to other (*ΔV*_self_ & *ΔV*_other_, respectively):

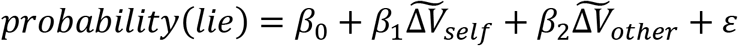

where probability to lie ranges from 0 to 1, and ~ represents the z-transformed monetary amounts. We refer to the estimated coefficients as *self-interest* (β*ΔV*_self_) and *regard for others* (β*ΔV*_other_).

#### Statistical analyses

All reported t-tests are two-tailed. All reported correlations are Pearson correlations.

### fMRI analysis

#### Statistical significance

For whole-brain analyses (*Neural correlates of dishonesty* and *Value to self and other* analyses), we used cluster-size threshold for multiple comparison correction. Cluster-defining threshold was set to 0.005, with 1000 Monte Carlo simulations to achieve a family-wise error of 0.05.

For ROI analyses (*Chosen value*, *Motive Modulation of value to self and other*, and *Motive modulation of the value system*) we used an FDR<0.05 threshold to correct for multiple comparisons.

#### Chosen value

To uncover voxels tracking the chosen value of each trial, we constructed a general linear model (GLM1) with the following predictors: (1) options period – a box-car function from trial onset until choice submission; (2) decision period – a box-car function from choice until ITI; and parametric modulators of the decision period: (3) chosen value – the amount of money (in ₪) that the Sender will receive in the chosen door; (4) unchosen value – the amount of money (in ₪) that the Sender would have received in the unchosen door; (5) Truth – an indicator function for trials in which the Sender told the truth; and (6) Lie – an indicator function for trials in which the Sender lied. All predictors were then convolved with HRF. Additional nuisance predictors included six motion-correction parameters and a mean signal from the ventricles, accounting for respiration. To reveal voxels tracking chosen value, we contrasted chosen value with unchosen value (predictors 3 and 4). We restricted the analysis to the valuation system only – the vmPFC and ventral striatum. We defined these areas using a mask generated from a meta-analysis of value studies^35^ (consisting of 385 voxels).

#### Neural correlates of dishonesty

We used GLM1 to contrast Truth (trials in which the participant sent a truthful message) and Lie trials (in which the participant sent a deceptive message) in a whole-brain analysis. The “chosen value” predictor controlled for the amount of money participants stood to gain on each trial, to ensure that the observed patterns of neural response reflected (dis)honesty, irrespective of reward size.

#### Value for self and other

To identify neural correlates of value to self (*ΔV*_self_) and value to other (*ΔV*_other_), we constructed a second general linear model (GLM2) with the following predictors: (1) options period – a box-car function from trial onset to choice submission; (2) decision period – a box-car function from choice until ITI; and parametric modulators of the options period: (3) *ΔV*_self_ – the difference between the profit to Sender (in ₪) in the Lie option and the Truth option ($Self_Lie_-$Self_Truth_); (4) *ΔV*_other_ – the difference between profit to Receiver (in ₪) in the Truth option and the Lie option ($Other_Truth_-$Other_Lie_). All predictors were then convolved with HRF. Additional nuisance predictors included six motion-correction parameters and a mean signal from the ventricles, accounting for respiration.

#### Motive modulation of value to self and other

To inspect the regions sensitive to value (to self or other) for individual differences in motives, we conducted ROI analyses. For the value to self (*ΔV*_self_) analysis, we focused on the LPFC and used an independently defined ROI^37^, a sphere consisting of 25 voxels around MNI coordinates - 48, 6, 28. We extracted the BOLD coefficients for *ΔV*_self_ (i.e., predictor 3 in GLM2) and correlated them with β*ΔV*_self_, estimated using participants’ behaviour. For the value to other (*ΔV*_other_) analysis, we used a 25-voxel sphere centred on MNI coordinates x, y, z: 54, −58, 22, based on previous studies^37,38^. We again extracted the BOLD signal, this time for *ΔV*_other_ (predictor 4 in GLM2) and correlated with behaviourally-estimated β*ΔV*_other_.

#### Motive modulation of the value system

To test how motives for dishonesty affect the valuation system, we started with computing for each participant an *other-self differential score* (β*ΔV*_other_–β*ΔV*_self_), indicating how much more one motive drives behaviour compared to the other. We then compared the neural activity tracking value to other with that of value to self (*ΔV*_other_ > *ΔV*_self_) in the valuation system. We defined the valuation system using a mask generated from a meta-analysis of value studies^35^ (consisting of 385 voxels). Finally, we regressed onto this map each participant’s other-self differential score, to identify motive-modulated value-representation. This analysis yielded a voxel-wise correlation map (correlating BOLD *ΔV*_other_ > *ΔV*_self_ and behavioural β*ΔV*_other_–β*ΔV*_self_), as implemented in BrainVoyager.

### Data Availability

All statistical maps and computer code used to analyse the fMRI data are available on OSF.org (https://osf.io/bvuxc/).

## Acknowledgements

We thank Liz Izakson for her help in data collection, and Dr. Xiaosi Gu for her thoughtful comments on the manuscript. This work was supported by National Science Foundation and US-Israeli Binational Science Foundation (NSF-BSF, grant #0612015334) and the Israel Science Foundation (ISF, grant #0612015023).

## Author contribution

DJL oversaw the project. AS and DJL designed the experiment. AS collected and analysed the data. AS and DJL wrote the manuscript.

## Additional Information

The authors declare no competing interests.

## Notes

https://osf.io/bvuxc/

## References

1. Gneezy, U. Deception: The role of consequences. American Economic Review 95, 384–394 (2005).

2. Mazar, N., Amir, O. & Ariely, D. The Dishonesty of Honest People: A Theory of Self-Concept Maintenance. Journal of Marketing Research 45, 633–644 (2008).

3. Jacobsen, C., Fosgaard, T. R. & Pascual-Ezama, D. Why do we lie? A practical guide to the dishonesy literature. Journal of Economic Surveys 32, 357–387 (2018).

4. Gneezy, U., Rockenbach, B. & Serra-Garcia, M. Measuring lying aversion. Journal of Economic Behavior and Organization 93, 293–300 (2013).

5. Pornpattananangkul, N., Zhen, S. & Yu, R. Common and distinct neural correlates of self-serving and prosocial dishonesty. Hum. Brain. Mapp. 39, 3086–3103 (2018).

6. Yin, L. & Weber, B. Can beneficial ends justify lying? Neural responses to the passive reception of lies and truth-telling with beneficial and harmful monetary outcomes. Soc Cogn Affect Neurosci 11, 423–432 (2016).

7. Cohn, A., Maréchal, M. A., Tannenbaum, D. & Zünd, C. L. Civic honesty around the globe. Science 365, 70–73 (2019).

8. Yin, L., Reuter, M. & Weber, B. Let the man choose what to do: Neural correlates of spontaneous lying and truth-telling. Brain and Cognition 102, 13–25 (2016).

9. Sip, K. E., Roepstorff, A., McGregor, W. & Frith, C. D. Detecting deception: the scope and limits. Trends in Cognitive Sciences 12, 48–53 (2008).

10. Jenkins, A. C., Zhu, L. & Hsu, M. Cognitive neuroscience of honesty and deception: a signaling framework. Current Opinion in Behavioral Sciences 11, 130–137 (2016).

11. Lisofsky, N., Kazzer, P., Heekeren, H. R. & Prehn, K. Investigating socio-cognitive processes in deception: A quantitative meta-analysis of neuroimaging studies. Neuropsychologia 61, 113–122 (2014).

12. Volz, K. G., Vogeley, K., Tittgemeyer, M., von Cramon, D. Y. & Sutter, M. The neural basis of deception in strategic interactions. Frontiers in Behavioral Neuroscience 9, (2015).

13. Spence, S. A. et al. Behavioural and functional anatomical correlates of deception in humans: Neuroreport 12, 2849–2853 (2001).

14. Abe, N. The neurobiology of deception: evidence from neuroimaging and loss-of-function studies. Current opinion in neurology 22, 594–600 (2009).

15. Christ, S. E., Essen, D. C. V., Watson, J. M., Brubaker, L. E. & Mcdermott, K. B. The Contributions of Prefrontal Cortex and Executive Control to Deception : Evidence from Activation Likelihood Estimate Meta-analyses. 1557–1566 (2009) doi:10.1093/cercor/bhn189.

16. Greene, J. D. & Paxton, J. M. Patterns of neural activity associated with honest and dishonest moral decisions. Proceedings of the National Academy of Sciences of the United States of America 106, 12506–12511 (2009).

17. Sip, K. E. et al. The production and detection of deception in an interactive game. Neuropsychologia 48, 3619–3626 (2010).

18. Abe, N. How the Brain Shapes Deception: An Integrated Review of the Literature. The Neuroscientist 17, 560–574 (2011).

19. Liang, C.-Y. et al. Neural correlates of feigned memory impairment are distinguishable from answering randomly and answering incorrectly: An fMRI and behavioral study. Brain and Cognition 79, 70–77 (2012).

20. Kozel, A. F. et al. A Pilot Study of Functional Magnetic Resonance Imaging Brain Correlates of Deception in Healthy Young Men. Journal Of Neuropsychiatry 16, 295–305 (2004).

21. Kozel, F. A. et al. Detecting Deception Using Functional Magnetic Resonance Imaging. (2005) doi:10.1016/j.biopsych.2005.07.040.

22. Langleben, D. D. et al. Telling truth from lie in individual subjects with fast event-related fMRI. Human Brain Mapping 26, 262–272 (2005).

23. Bhatt, S. et al. Lying about facial recognition: An fMRI study. Brain and Cognition 69, 382–390 (2009).

24. Rilling, J. K. et al. A Neural Basis for Social Cooperation. Neuron 35, 395–405 (2002).

25. Andreoni, J. & Miller, J. Giving According to GARP: An Experimental Test of the Consistency of Preferences for Altruism. Econometrica 70, 737–753 (2002).

26. Moll, J. et al. Human fronto-mesolimbic networks guide decisions about charitable donation. Proceedings of the National Academy of Sciences 103, 15623–15628 (2006).

27. Hare, T. a, Camerer, C. F., Knoepfle, D. T. & Rangel, A. Value computations in ventral medial prefrontal cortex during charitable decision making incorporate input from regions involved in social cognition. The Journal of neuroscience: the official journal of the Society for Neuroscience 30, 583–590 (2010).

28. Zaki, J. & Mitchell, J. P. Equitable decision making is associated with neural markers of intrinsic value. Proceedings of the National Academy of Sciences 108, 19761–19766 (2011).

29. Fehr, E. & Camerer, C. F. Social neuroeconomics: the neural circuitry of social preferences. Trends in Cognitive Sciences 11, 419–427 (2007).

30. Ruff, C. C. & Fehr, E. The neurobiology of rewards and values in social decision making. Nature reviews. Neuroscience 15, 549–562 (2014).

31. Janowski, V., Camerer, C. & Rangel, A. Empathic choice involves vmPFC value signals that are modulated by social processing implemented in IPL. Social Cognitive and Affective Neuroscience 8, 201–208 (2013).

32. Zaki, J., Lopez, G. & Mitchell, J. P. Activity in ventromedial prefrontal cortex co-varies with revealed social preferences: evidence for person-invariant value. Social Cognitive and Affective Neuroscience 9, 464–469 (2014).

33. Morelli, S. A., Sacchet, M. D. & Zaki, J. Common and distinct neural correlates of personal and vicarious reward: A quantitative meta-analysis. NeuroImage 112, 244–253 (2015).

34. Smith, D. V., Clithero, J. A., Boltuck, S. E. & Huettel, S. A. Functional connectivity with ventromedial prefrontal cortex reflects subjective value for social rewards. Social Cognitive and Affective Neuroscience 9, 2017–2025 (2013).

35. Bartra, O., McGuire, J. T. & Kable, J. W. The valuation system: A coordinate-based meta-analysis of BOLD fMRI experiments examining neural correlates of subjective value. NeuroImage 76, 412–427 (2013).

36. Levy, D. J. & Glimcher, P. W. The root of all value: a neural common currency for choice. Current Opinion in Neurobiology 1–12 (2012) doi:10.1016/j.conb.2012.06.001.

37. Crockett, M. J., Siegel, J. Z., Kurth-Nelson, Z., Dayan, P. & Dolan, R. J. Moral transgressions corrupt neural representations of value. Nature Neuroscience 20, 879–885 (2017).

38. Bzdok, D. et al. Parsing the neural correlates of moral cognition: ALE meta-analysis on morality, theory of mind, and empathy. Brain Structure and Function 217, 783–796 (2012).

39. Erat, S. & Gneezy, U. White Lies. Management Science 58, 723–733 (2012).

40. Milosavljevic, M., Malmaud, J., Huth, A., Koch, C. & Rangel, A. The Drift Diffusion Model Can Account for the Accuracy and Reaction Time of Value-Based Choices Under High and Low Time Pressure. SSRN Electronic Journal (2010) doi:10.2139/ssrn.1901533.

41. Krajbich, I., Bartling, B., Hare, T. & Fehr, E. Rethinking fast and slow based on a critique of reaction-time reverse inference. Nature Communications 6, 1–9 (2015).

42. Krajbich, I., Lu, D., Camerer, C. & Rangel, A. The Attentional Drift-Diffusion Model Extends to Simple Purchasing Decisions. Frontiers in Psychology 3, (2012).

43. Carlson, R. W. & Crockett, M. J. The lateral prefrontal cortex and moral goal pursuit. Current Opinion in Psychology 24, 77–82 (2018).

44. Zhu, L. et al. Damage to dorsolateral prefrontal cortex affects tradeoffs between honesty and self-interest. Nature neuroscience 17, 1319–21 (2014).

45. Maréchal, M. A., Cohn, A., Ugazio, G. & Ruff, C. C. Increasing honesty in humans with noninvasive brain stimulation. Proceedings of the National Academy of Sciences 114, 4360–4364 (2017).

46. Carter, R. M., Bowling, D. L., Reeck, C. & Huettel, S. A. A Distinct Role of the Temporal-Parietal Junction in Predicting Socially Guided Decisions. Science 337, 109–111 (2012).

47. Saxe, R. & Kanwisher, N. People thinking about thinking peopleThe role of the temporo-parietal junction in “theory of mind”. NeuroImage 19, 1835–1842 (2003).

48. Samson, D., Apperly, I. A., Chiavarino, C. & Humphreys, G. W. Left temporoparietal junction is necessary for representing someone else’s belief. Nature Neuroscience 7, 499–500 (2004).

49. Schurz, M., Radua, J., Aichhorn, M., Richlan, F. & Perner, J. Fractionating theory of mind: A meta-analysis of functional brain imaging studies. Neuroscience & Biobehavioral Reviews 42, 9–34 (2014).

50. Young, L., Cushman, F., Hauser, M. & Saxe, R. The neural basis of the interaction between theory of mind and moral judgment. Proceedings of the National Academy of Sciences 104, 8235–8240 (2007).

51. Young, L., Camprodon, J. A., Hauser, M., Pascual-Leone, A. & Saxe, R. Disruption of the right temporoparietal junction with transcranial magnetic stimulation reduces the role of beliefs in moral judgments. Proceedings of the National Academy of Sciences 107, 6753–6758 (2010).

52. Clithero, J. a & Rangel, A. Informatic parcellation of the network involved in the computation of subjective value. Social Cognitive and Affective Neuroscience (2013) doi:10.1093/scan/nst106.

