## Supplementary material for "Contribution of self- and other-regarding motives to (dis)honesty": Table S1 and Figure S1-S3

**Table S1. List of payoffs on each trial.** $Self Lie and $Other Lie are payoffs to the Sender and Receiver, respectively, if the Sender chooses to Lie. $Self Truth and $Other Truth are payoffs to the Sender and Receiver, respectively, if the Sender chooses to tell the truth. Catch trials, in which the Sender's payoff is the same for both alternatives, are marked with an asterisk.

| | **$Self Lie** | **$Other Lie** | **$Self Truth** | **$Other Truth** |
|---|---|---|---|---|
| | 21 | 7 | 18 | 12 |
| | 36 | 10 | 34 | 30 |
| | 26 | 3 | 23 | 8 |
| | 35 | 2 | 32 | 22 |
| | 24 | 4 | 19 | 14 |
| | 30 | 13 | 29 | 26 |
| | 35 | 13 | 32 | 31 |
| | 26 | 6 | 24 | 19 |
| | 28 | 9 | 23 | 10 |
| | 35 | 2 | 32 | 8 |
| | 17 | 3 | 14 | 11 |
| | 29 | 10 | 27 | 17 |
| | 15 | 3 | 10 | 9 |
| | 22 | 7 | 18 | 13 |
| | *17 | 4 | 17 | 6 |
| | 19 | 5 | 14 | 8 |
| | 15 | 1 | 10 | 3 |
| | 15 | 4 | 11 | 8 |
| | 32 | 11 | 28 | 18 |
| | 21 | 9 | 18 | 12 |
| | 31 | 9 | 19 | 16 |
| | 27 | 5 | 17 | 14 |
| | 36 | 23 | 29 | 27 |
| | 33 | 6 | 27 | 9 |
| | 28 | 1 | 17 | 5 |
| | 23 | 17 | 19 | 18 |
| | 28 | 12 | 25 | 24 |
| | 24 | 8 | 12 | 10 |
| | 42 | 18 | 33 | 26 |
| | 19 | 4 | 11 | 7 |
| | 19 | 3 | 17 | 8 |
| | 24 | 9 | 21 | 10 |
| | 35 | 14 | 33 | 26 |
| | 16 | 1 | 12 | 5 |
| | 16 | 3 | 11 | 4 |
| | 33 | 12 | 28 | 27 |
| | *38 | 15 | 38 | 23 |
| | 34 | 13 | 33 | 22 |
| | 38 | 8 | 33 | 19 |
| | 35 | 13 | 30 | 23 |
| **mean** | **26.925** | **8** | **22.45** | **15.2** |
| **SD** | **7.74** | **5.26** | **8.21** | **8.06** |
| **minimum** | **15** | **1** | **10** | **3** |
| **maximum** | **42** | **23** | **38** | **31** |

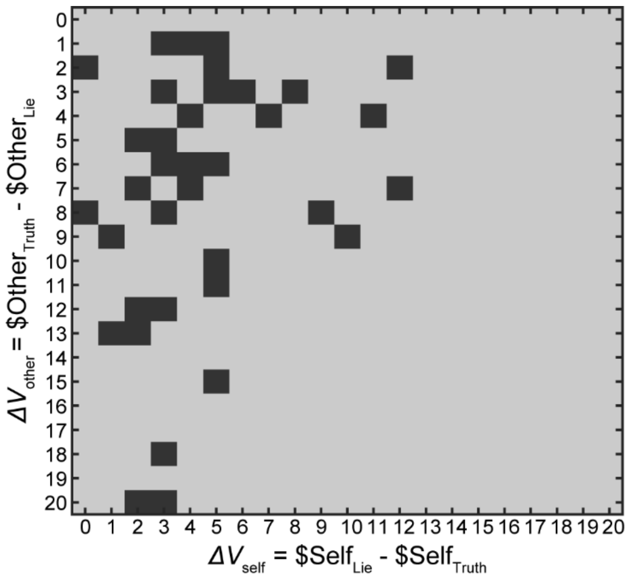


**Figure S1. Self and Other payoffs.** Potential losses to the Receiver (value to other) on the y-axis and potential gains for the Sender (value to self) on the x-axis. Each dark square represents a unique trial in our experiment.

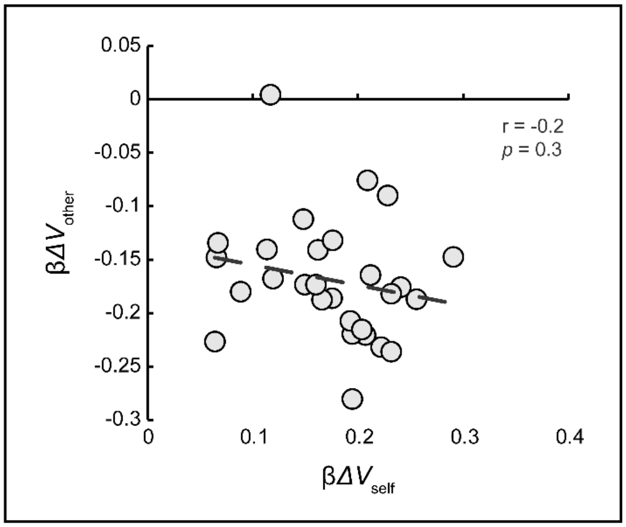


**Figure S2. Self-interest and regard for others are not correlated.** Coefficient of potential losses to the Receiver (value to other) on the y-axis, and coefficient of potential gains for the Sender (value to self) on the x-axis. Each circle represents a subject. N=28.

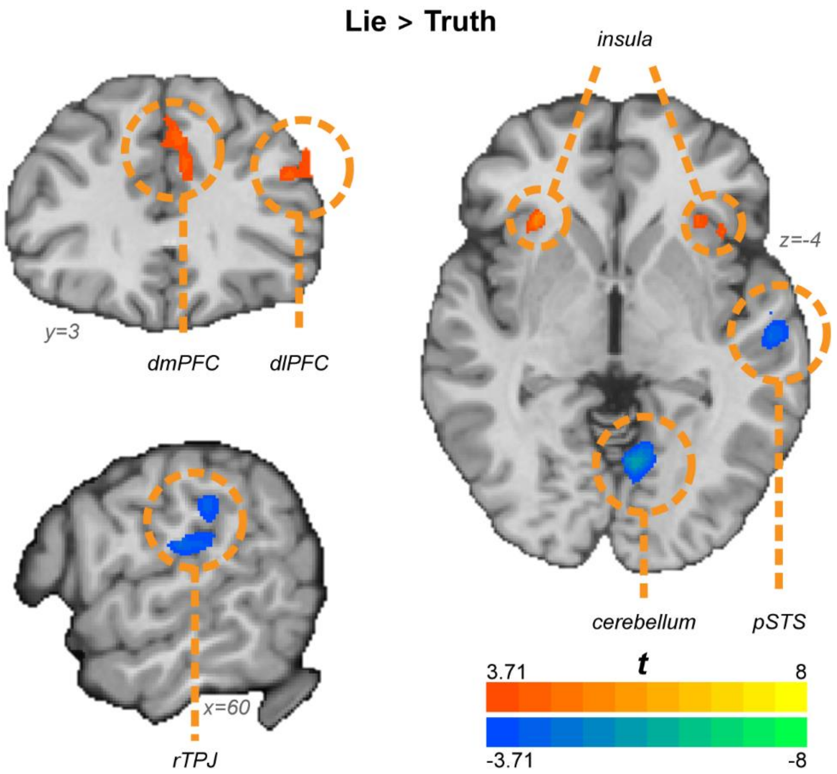


**Figure S3. Lie vs. Truth brain map.** Contrasting trials in which subjects chose Deceitful message with trials in which they chose the Truthful message. Map thresholded at $p$=0.001, cluster-size corrected. N=27.

### Supplementary analysis & results:

A potential explanation to negative relationship between value to self and the LPFC comes from the role of the LPFC in cognitive control (Badre and Nee 2018) – participants might experience more conflict and need for control when the potential reward from lying ($\$Self_{Lie}$-$\$Self_{Truth}$) is small. If this was the case, we would expect to find a negative relationship between the potential reward from lying and reaction times, such that smaller differences would yield longer reaction times. Because our neural model controls for the effects of reaction times on value representation, it is a less plausible explanation for the observed result. Nonetheless, we directly tested this hypothesis.

To examine how potential profits from lying affect decision time, we conducted two analyses. First, we regressed value to self ($\Delta V_{self}$) onto reaction times, clustering the errors per participant, to get an across-participant measure of the relationship between the two variables. Second, we ran participant-specific correlations of reaction times and value to self, yielding a correlation coefficient per participant. Then, we examined whether this resulting correlation is related to other behavioral measures by correlating it with the overall dishonesty and with self-regarding motive for dishonesty.

We find no significant relationship between reaction times and $\Delta V_{self}$ ($\beta\Delta V_{self}$=0.005, $p$=0.48). Furthermore, each participant's correlation coefficient between reaction times and $\Delta V_{self}$ is unrelated to their $\beta\Delta V_{self}$ (r(27)=-0.02, $p$=0.89). That is, the relationship between potential profits and reaction times does not predict levels of self-interest. Interestingly, it is negatively linked to overall dishonesty (r(27)=-0.74, $p$<0.001), such that honest participants take longer to choose when they can gain a large profit from lying, while dishonest participants take longer to choose when the potential profit from lying is small.
